# Fetal lung vascular development is disrupted by mechanical compression and rescued by administration of amniotic fluid stem cell extracellular vesicles via regulation of the Hippo signaling pathway

**DOI:** 10.1101/2023.11.30.569427

**Authors:** Rebeca L. Figueira, Kasra Khalaj, Lina Antounians, Fabian Doktor, Maria Sole Gaffi, Mikal Obed, Sree Gandhi, Augusto Zani

## Abstract

Postnatal pulmonary hypertension is the biggest treatment challenge and major determinant for poor outcome in infants with congenital diaphragmatic hernia (CDH). CDH lungs are hypoplastic and exhibit vascular remodeling, whose pathogenesis remains poorly understood. Using a novel micro-static compression system, herein we found that mechanical compression induces vascular remodeling and downregulation of key angiogenic markers in rat and human fetal lung models, with similar features observed in CDH fetal lung autopsy samples. These fetal lung vascular changes are reversed back to normal upon administration of extracellular vesicles derived from amniotic fluid stem cells (AFSC-EVs), a regenerative approach previously shown to restore lung branching morphogenesis and epithelial differentiation in CDH models. Exploring pathways that are dysregulated in CDH lungs and involved in mechanotransduction, we found that compressed fetal lungs had altered expression of Hippo signaling factors that was restored upon AFSC-EV administration. We found that AFSC-EV cargo contains some miRNAs involved in lung vascular development and Hippo pathway, indicating that AFSC-EV regenerative effects is associated with the delivery of specific miRNAs. This study uncovers the role of mechanical compression that herniated organs exert on CDH fetal lungs and proposes a new cell-free strategy to restore normal fetal lung vascular development.

## Introduction

Normal pulmonary vascular development is a critical factor for the cardiorespiratory adaptation of the newborn ^1^. Disruption of the vascular network in fetal lungs is a primary cause for postnatal pulmonary hypertension, a determinant for the poor outcome of affected babies ^2,3^. There is a wide range of conditions affecting the mother or the fetus that can cause abnormal pulmonary vascular development leading to postnatal pulmonary hypertension ^4-7^. One of these is congenital diaphragmatic hernia (CDH), a complex birth defect characterized by an incomplete formation of the diaphragm and herniation of abdominal viscera into the chest ^8^. CDH lungs are hypoplastic, with disrupted branching morphogenesis (impaired growth), arrested epithelial and mesenchymal cell differentiation (impaired maturation), and vascular remodeling with fewer vessels that have structural alterations of their wall composition (impaired vascularization) ^8-11^. Particularly, the hypermuscularization of the preacinar arteries is associated with impaired relaxation of pulmonary arteries leading to high vascular resistance, which manifests postnatally as pulmonary hypertension. Pulmonary hypertension is a primary determinant for the high mortality and morbidity in infants with CDH, as its management remains one of the greatest challenges of neonatal medicine ^12,13^. The challenge is twofold: the incomplete understanding of the pathophysiology of antenatal vascular remodeling in CDH lungs and the lack of a therapy that would promote normal vascular development thus preventing the occurrence of pulmonary hypertension.

Although the pathophysiology of pulmonary hypoplasia secondary to CDH has been studied for decades, the role of mechanical compression exerted by herniated organs onto the fetal lungs remains incompletely understood. Several experimental studies have shown that mechanical compression on fetal lungs induces an arrest in branching morphogenesis and an impairment in lung maturation ^14-16^. However, the effects of mechanical compression on lung vascular development in isolation have not been characterized. In the current study, using a novel micro-static compression system, we found that mechanical compression induces vascular remodeling and downregulation of key angiogenic markers in rat and human fetal lung models, with similar features observed in CDH fetal lung autopsy samples. Exploring pathways that are dysregulated in CDH lungs and involved in mechanotransduction, we found that compressed fetal lungs have altered expression of Hippo signaling factors.

It is recognized that current therapies for pulmonary hypertension are not effective in all cases, likely as they do not address the underlying vascular remodeling that is inherent to CDH hypoplastic lungs. In general, there is consensus that rescuing lung development *in utero* is the ideal approach to treat hypoplastic lungs in babies with CDH ^8,17^. Fetoscopic endoluminal occlusion (FETO), a non-pharmacological approach that improves survival in selected cases of severe CDH by improving fetal lung growth and maturation, does not rescue fetal lung vascular development ^17-19^. Ideally, an optimal therapy would be one that addresses all aspects of fetal lung underdevelopment, that is lung growth, maturation, and vascularization. Experimentally, administration of extracellular vesicles derived from amniotic fluid stem cells (AFSC-EVs) has been reported to rescue branching morphogenesis and epithelial and mesenchymal cell differentiation in rodent and human models of CDH ^20-22^. The beneficial effect of AFSC-EVs on hypoplastic lungs is attributed at least in part to the delivery of their microRNA cargo ^20,21^. In the current study, we show that AFSC-EV administration to a model of fetal lung mechanical compression reverses vessel density, medial wall thickness, and expression of markers involved in angiogenesis and Hippo signaling pathway back to normal. We report that AFSC-EV cargo is enriched in miRNAs that regulate angiogenesis and/or vasculogenesis, as well as processes involved in the Hippo signaling pathway.

Taken together, the findings reported in this study uncover the role of mechanical compression that herniated organs exert on CDH fetal lungs, thus increasing our knowledge on the pathophysiology of vascular remodeling in pulmonary hypoplasia. Moreover, this study adds to the growing evidence that an EV-based therapy could restore normal fetal lung development in all its aspects, including vessel development.

## RESULTS

### Mechanical compression disrupts fetal vascular development in fetal rat lung explants

To explore the effects of mechanical compression on lung vascular development, we established a novel *ex vivo* model of CDH by creating a static micro-compression system that simulates the herniation of the abdominal organs into the thoracic cavity, as it occurs in human fetuses with CDH (**Figure 1A-B**). We harvested lungs at embryonic day (E) 19.5 from fetal rats of either olive oil-gavaged dams or nitrofen-gavaged dams. The nitrofen model of CDH is the most widely used and is based on maternal administration of nitrofen at E9.5, which causes a diaphragmatic defect in about half of the litter and pulmonary hypoplasia in all fetuses ^23,24^. For these experiments, we exclusively selected nitrofen-exposed lungs from fetuses that did not develop a diaphragmatic defect in order to avoid the confounding effect of *in utero* compression. Normal and nitrofen-exposed lungs were either subjected to compression (normal+compression and nitrofen+compression groups) or grown without compression (normal and nitrofen groups). This model gave us the opportunity to evaluate the role of mechanical forces on normal fetal lung development. We first verified the effects of mechanical compression on lung growth, measured as surface area and number of airspaces. Compared to compressed and non-compressed normal lungs nitrofen-exposed lungs had a reduced surface area and number of airspaces regardless of whether compression was applied, confirming the detrimental effect of nitrofen on branching morphogenesis, as previously reported (**Suppl.Fig. 1A-B**) ^15^. Normal compressed lungs had a reduced surface area and number of airspaces compared to non-compressed normal lungs, showing that compression alone affects branching morphogenesis (**Suppl.Fig. 1A-B**) ^14-15^. We then examined the effects of compression in the pulmonary vascular development. We observed that compared to non-compressed lungs, normal and nitrofen compressed fetal lung explants had decreased vascular density, increased medial wall thickness of the pulmonary arteries, and downregulated gene expression of angiogenic factors, such as vascular endothelial growth factor-a (*Vegfa)*, vascular endothelial growth factor receptor 2 (*Vegfr2)*, platelet endothelial cell adhesion molecule (*Cd31)*, endothelial nitric oxide synthase (*Enos)*, endothelial PAS domain-containing protein 1 (*Epas1)*, endothelial transcription factor gata-2 (*Gata2)*, angiopoietin 1 (*Ang1),* and angiopoietin 1 receptor (*Tie2)* (**Figure 1C-E**). Moreover, normal and nitrofen compressed lung explants had reduced protein expression of VEGFA, vascular endothelial growth factor receptor 1 (VEGFR1), VEGFR2, CD31 and ENOS in comparison to non-compressed lungs (**Figure 1F**).

**Figure 1.**
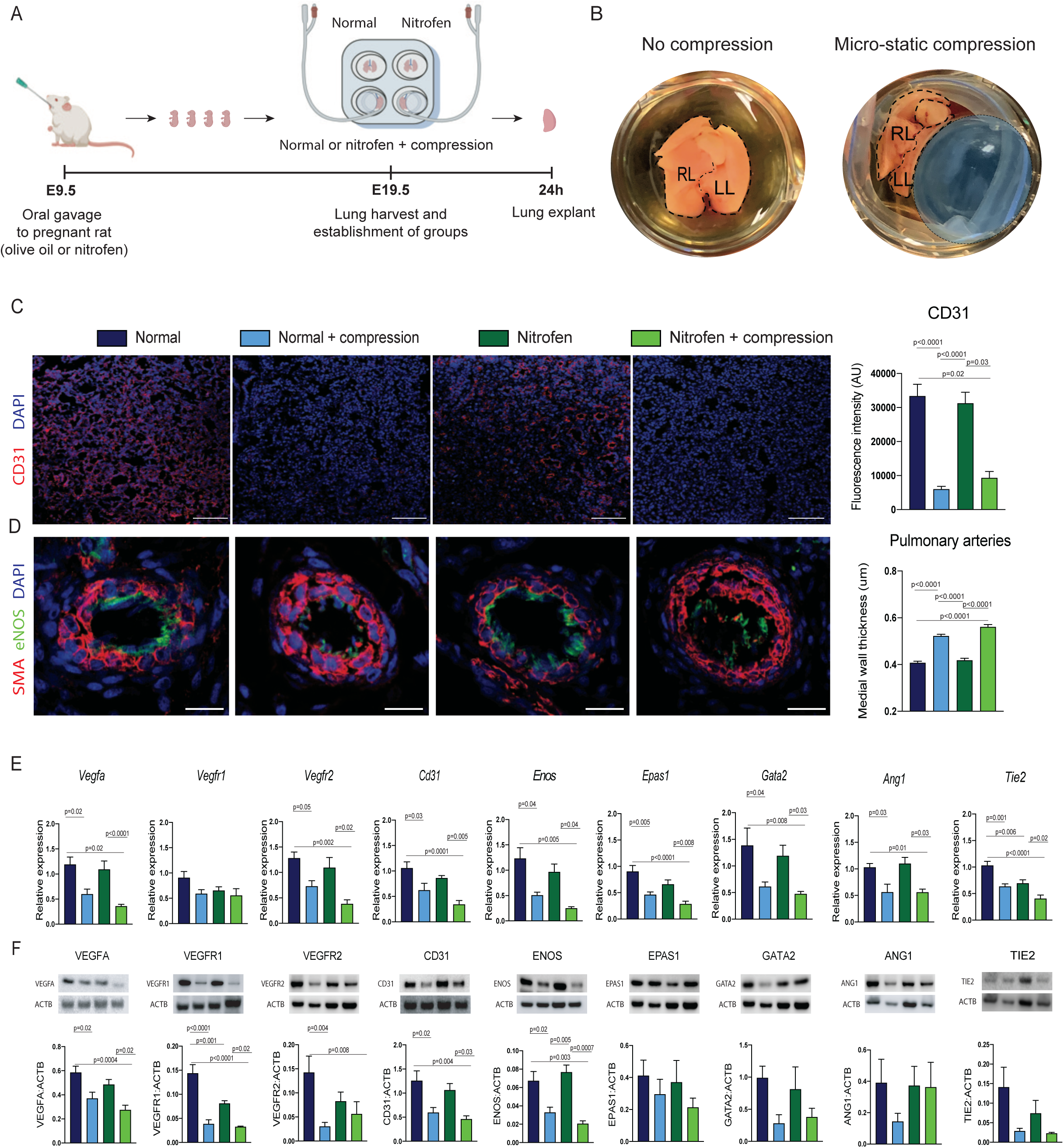
Mechanical compression impairs vascular development in both normal and hypoplastic rat fetal lungs in an ex-vivo CDH model. (A) Timeline of *ex vivo* studies in the rat model. (B) Representative macroscopic photos of fetal lungs not subjected to compression (normal and nitrofen groups), and of fetal lungs that were compressed using a static micro-compression system (normal + compression and nitrofen + compression groups). RL: right lung; LL: left lung. (C) Representative immunofluorescence images of the pan-endothelial marker CD31 (red) and nuclear marker 4′,6-diamidino-2-phenylindole (DAPI, blue) in rat fetal lung explants from four experimental groups (n=5 in each group). Group comparison of CD31 fluorescence intensity (Arbitrary Units, AU; 5 fields per biological replicate). Scale bar = 50 µm. (D) Representative immunofluorescence images of pulmonary arteries co-stained with smooth muscle actin (SMA; red), endothelial nitric oxide synthase (eNOS; green), and nuclear marker (DAPI, blue). Medial wall thickness of pulmonary arteries from Normal (n=6), Normal + compression (n=8), Nitrofen (n=9) and Nitrofen + compression (n=10) groups (10 fields per sample containing a minimum of 3 pulmonary arteries each; µm). Scale bar = 50 µm. (E) Group comparison of gene expression levels of angiogenic factors, such as vascular endothelial growth factor A (*Vegfa*) and its receptors *Vegfr1* and *Vegfr2*; platelet endothelial cell adhesion molecule (*Cd31*), endothelial nitric oxide synthase (*Enos*); endothelial PAS domain protein 1 (*Epas1*); endothelial transcription factor (*Gata2*), angiopoietin 1 (*Ang1*) and its receptor TEK tyrosine kinase (*Tie2*) relative to glyceraldehyde-3-phosphate dehydrogenase (*Gapdh*) expression (n=5 in each group). (F) Representative Western blotting images and quantification of the same angiogenic factors as in (E) relative to actin-b (ACTB), (n=4 in each group). (A-F) Data are shown as mean ± standard deviation (SD). Groups were compared using Kruskal-Wallis (post hoc Dunn’s nonparametric comparison) for [C; H = 48.03], (D; H value = 155.4) and (E; *Vegfa,* F = 7.169 and *Ang1,* H = 16.47 and F; EPAS1, H = 2.75; and TIE2, H = 9.90); and one-way ANOVA (Tukey post-test) for (E; *Vegfr1,* F = 2.245; *Vegfr2,* F = 6.68; *Cd31,* F = 9.41*; Enos,* F = 6.54; *Epas1,* F = 11.6; *Gata2*, F = 6.53; and *Tie2,* F = 15.22) and (F; VEGFA, F = 7.55; VEGFR1, F = 22.48; VEGFR2, F = 16.75; CD31, F = 6.55; ENOS, F = 13.89; GATA2, F = 7.20; ANG1; and F = 4.39), according to Shapiro-Wilk normality test. Only p-values < 0.05 are shown.

### Vascular development is impaired in an *ex-vivo* human fetal lung model of mechanical compression

Next, we sought to investigate whether the effects of mechanical compression observed in our model of rat fetal lung explants could be recapitulated in the human lung. To this end, we established an *ex vivo* human fetal lung model, where lung specimens were obtained from terminations of healthy fetuses conducted at 18–19 weeks of gestation (canalicular stage of lung development) (**Figure 2A**). Explants were either compressed using our compression model for 24h, or not compressed. First, we evaluated lung growth and morphogenesis using radial airspace count on compressed and non-compressed fetal human lungs. We observed that compared to non-compressed explants, human fetal lung specimens subjected to mechanical compression had fewer airspaces (**Figure 2B)**. To validate our model, we then compared compressed fetal lung explants to human lung autopsy of fetuses with CDH and compared them to human lung autopsy of sex and age-matched controls. Compressed fetal human lungs had decreased number of airspaces at similar levels to those observed in CDH lung autopsy samples (**Figure 2B**). We then investigated whether mechanical compression of human fetal lung explants would also impair vascular development. Compared to non-compressed lung explants, compressed human fetal lungs had decreased vascular density and increased medial wall thickness of the pulmonary arteries (**Figure 2C-D**). These features replicated the vascular phenotype of CDH autopsy lungs (**Figure 2C-D**). Moreover, compared to non-compressed human fetal lung explants, compressed explants had downregulated gene expression of *VEGFA*, *VEGFR1*, *VEGFR2*, *CD31* and *ENOS* and decreased protein expression of VEGFR1, CD31, ENOS, EPAS, GATA2 and TIE2 (**Figure 2E-F**).

**Figure 2.**
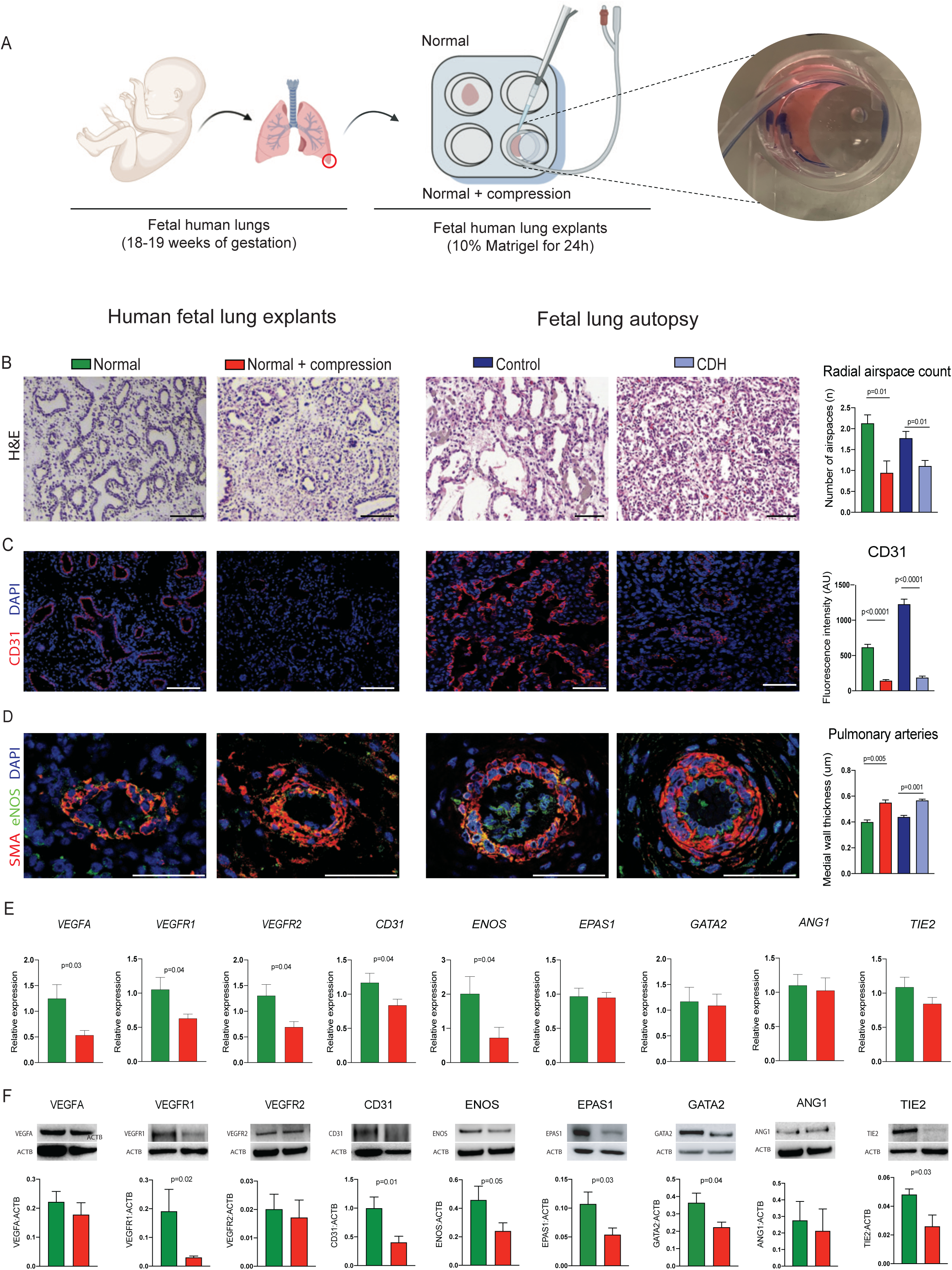
Mechanical compression alone disrupts vascular development in human lungs. (A) Experimental design and representative images of human fetal lung explants not subjected to compression (normal group) or compressed using a static micro-compression system (normal + compression group). (B) Representative histology images (hematoxylin/eosin staining) of human fetal lung explants (normal and normal + compression groups) and human fetal lung autopsy samples of healthy fetuses (controls with no pulmonary disease) and fetuses with CDH. Group comparison of radial airspace count (RAC) to assess severity of pulmonary hypoplasia in human fetal lung explants from Normal (n=5), Normal + compression (n=6), and human fetal lung autopsy samples from Control (n=4) and CDH (n=4). Scale bar = 50 µm. (C) Representative immunofluorescence images of CD31 in human fetal lung explants (n = 4 in each group) and human fetal lung autopsy samples. Scale bar = 50 µm. Group comparison of CD31 fluorescence intensity (AU; 10 fields per biological replicate). (D) Representative immunofluorescence images of pulmonary arteries co-stained with SMA (red), eNOS (green), and DAPI (blue) in human fetal lung explants (n = 4 in each group) and human fetal lung autopsy samples. Scale bar = 50 µm. Group comparison for medial wall thickness (10 fields per sample containing a minimum of 3 pulmonary arteries each). (E) Group comparison of gene expression of angiogenic factors described in Fig.1E relative to GAPDH in human fetal lung explants (n = 4 in each group). (F) Representative Western blotting images and quantification of angiogenic factors relative to ACTB in human fetal lung explants (n = 8 in each group). (A-F) Data are shown as mean ± SD. Groups were compared using Kruskal-Wallis (post hoc Dunn’s nonparametric comparison) for (B; H = 16.03) and (C; H = 103.9); one-way ANOVA (Tukey post-test) for (D; F = 20.13); two-tailed Student’s t-test for (E; *VEGFA,* t = 2.36/ df = 13; *VEGFR1,* t = 2.27/ df = 10; *VEGFR2*, t = 2.57/ df 6; *CD31,* t = 2.07/ df = 30; TIE2, t = 1.46/ df = 20; and *ANG1,* t = 0.29/df = 22) and (F; VEGFA, t = 0.81/df = 20; CD31, t = 2.76/ df = 19; EPAS1, t = 2.21/ df = 24; TIE2, t = 2.44/ df = 17), and two-tailed Mann Whitney test for (E; *EPAS1,* U = 427; *ENOS,* U = 14; *GATA2,* U = 88) and (F; VEGFR1, U = 22; VEGFR2, U = 26; ENOS, U = 59; GATA2, U = 37; ANG1, U = 63), according to Shapiro-Wilk normality test.

### Mechanical compression disrupts the hippo signaling pathway in rat and human fetal lungs

Next, we sought to explore the underlying mechanisms of compression-induced vascular remodeling in fetal lungs. Using a dataset that we previously obtained by single nucleus RNA-sequencing of rat fetal lungs with CDH ^25^, we investigated possible signaling networks that are known to be associated with mechanical stimuli response. We found that CDH lungs had dysregulated expression of Hippo pathway genes across the main cell populations of the fetal lung, i.e., endothelium, epithelium, mesenchyme, and immune cells (**Figure 3A-C**). The Hippo pathway is a conserved signaling pathway well known by its mechanosensitive regulators that control important biological processes involved in cell proliferation, survival, and differentiation^26-27^. When we analyzed the Hippo pathway genes dysregulated in rat CDH lungs, we found that that normal rat lung explants subjected to compression had downregulation of serine/threonine kinase 3 and 4 (*Stk3, Stk4),* large tumor suppressor kinase 1/2 (*Lats1/2*), MOB kinase activator 1A/B *(Mob1a, Mob1b),* yes-associated protein 1 (*Yap),* WW-domain containing transcriptional regulator 1 (*Taz),* transcriptional enhanced associate domain (*Tead),* cysteine-rich angiogenic inducer 61 *(Cyr61),* connective tissue growth factor (*Ctgf),* and ankyrin repeat domain 1 (*Ankrd1)* in comparison to non-compressed normal lung explants (**Figure 3D**). Similarly, nitrofen compressed lungs had downregulated *Stk3, Lats2, Mob1a, Mob1b, Yap, Taz, Tead, Cyr61, Ctgf,* and *Ankrd1* genes when compared to non-compressed nitrofen lung explants (**Figure 3D**). In our *ex vivo* human model of CDH, we observed that compressed human fetal lung explants had downregulated gene expression of *STK3,* salvador1 (*SAV1), YAP, TAZ, TEAD, CYR61, CTGF* and *ANKRD1* (**Figure 3E**).

**Figure 3.**
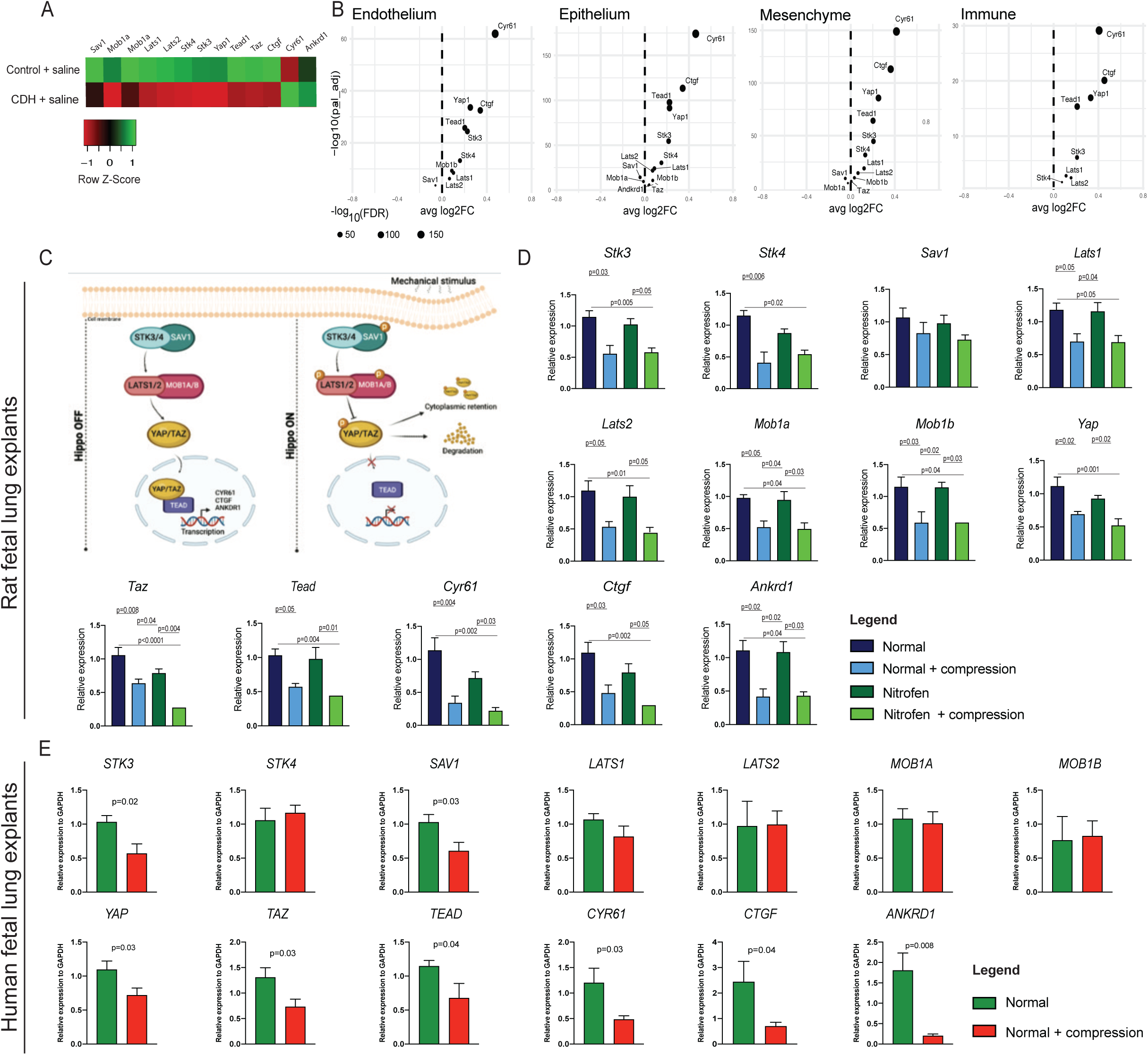
Mechanically compressed rat and human lungs have downregulated expression of Hippo signaling pathway genes. (A) Heatmap of differentially expressed Hippo signaling factors in Control + saline and CDH + saline rat fetal lungs. P-value adjusted < 0.05 for each gene. (B) Volcano plots showing average log2 fold change (FC) of Hippo signaling pathway factors in Control + saline and CDH + saline rat fetal lungs by major cell type (endothelium, epithelium, immune, and mesenchyme). The size of the node is proportional to -log10 (false discover rate; FDR). Log2(FC) >0 on the right side of the plot indicates higher expression in Control + saline vs. CDH + saline rat fetal lungs. (C) Graphical representation of the Hippo pathway cascade in mammals. The left side of panel represents the pathway when inactive, i.e. OFF; the right side of the panel represents the pathway when activated by a mechanical stimulus, i.e. ON. (D) Group comparison of gene expression of upstream transcription factors (*Stk3, Sav1, Yap, Taz* and *Tead*) and downstream effectors (*Cyr61, Ctgf*, and *Ankdr1*) of the Hippo pathway in fetal rat lung explants (n = 5 in each group). (E) Group comparison of gene expression of Hippo pathway upstream and downstream factors in human lung explants (n = 5) in each group. (D-E) Data are shown as mean ± SD. Rat fetal lungs were compared using one-way ANOVA (Tukey post-test) for (D; Sav1, F = 1.19*; Lats1,* F = 4.91; *Lats2,* F = 14.38*; Mob1a,* F = 5.62*; Mob1b,* F = 5.96*; Yap,* F = 7.10*; Taz,* F = 10.75; and *Tead,* F = 6.81) and Kruskal-Wallis (post hoc Dunn’s nonparametric comparison) for (D; *Stk3,* U *=* 15.57*; Stk4,* U = 13.32; *Cyr6,* U = 19.52*; Ctgf,* U = 16.07; and *Ankdr1,* U = 16.61) according to Shapiro-Wilk normality test. Human fetal lung explants were compared using two-tailed Student’s t-test for (E; *STK3,* t = 2.78/ df = 13*; SAV1,* t = 2.49/ df = 11; *LATS1,* t = 1.42/ df = 28; *MOB1A,* t = 0.28/ df = 17; *MOB1B,* t = 0.44/ df = 15; *YAP,* t = 2.20/ df = 20; *TAZ,* t = 2.38/ df = 15; *TEAD,* t = 2.25/ df = 12; *CYR61*, t = 2.41/ df = 12; *CTGF,* t = 2.33/ df = 9; and *ANKDR1,* t = 3.38/ df = 9) and two-tailed Mann Whitney test for (E; *STK4,* U = 10; and *LATS2,* U = 11), according to Shapiro-Wilk normality test. Only p-values < 0.05 are shown.

### Impaired vascular development is rescued by administration of AFSC-EVs to fetal lung explants

To investigate whether the administration of AFSC-EVs could rescue fetal lung vascular development in mechanically compressed fetal lungs, we injected rat AFSC-EVs (rAFSC-EVs) or human AFSC-EVs (hAFSC-EVs) in our established *ex vivo* rat and human fetal models of mechanical compression, respectively rAFSC-EVs and hAFSC-EVs were characterized as shown in **Suppl.Fig.2**. We first evaluated whether intratracheal administration of rAFSC-EVs would rescue lung branching morphogenesis using our *ex vivo* model of compression in rats (**Figure 4A**). We observed that compared to untreated lungs, mechanically compressed lungs treated with rAFSC-EVs had similar number of airspaces to non-compressed normal lung explants (**Suppl.Fig. 1C**). We also observed that intratracheal administration of rAFSC-EVs rescued vascular density, restored the medial wall thickness of the pulmonary arteries, as well as the gene expression of key angiogenic markers back to normal levels (*Vegfr2, Cd31, eNOS, Gata2* and *Ang1)* (**Figure 4B-D**). We then investigated whether the administration of hAFSC-EVs could also improve vascular development in human specimens using our *ex vivo* model of compression in human fetal lungs (**Figure 4E**). We first confirmed that the topical administration of hAFSC-EVs to mechanically compressed human fetal lung explants rescued the number of airspaces back to normal levels (**Suppl.Fig. 1D**). Administration of hAFSC-EVs to compressed human fetal lungs also rescued vascular density, reduced the medial wall thickness of the pulmonary arteries, and restored gene expression of *VEGFA, VEGFR2, CD31, EPAS1, TIE2* and *ANG1* **(Figure 4F-H)**.

**Figure 4.**
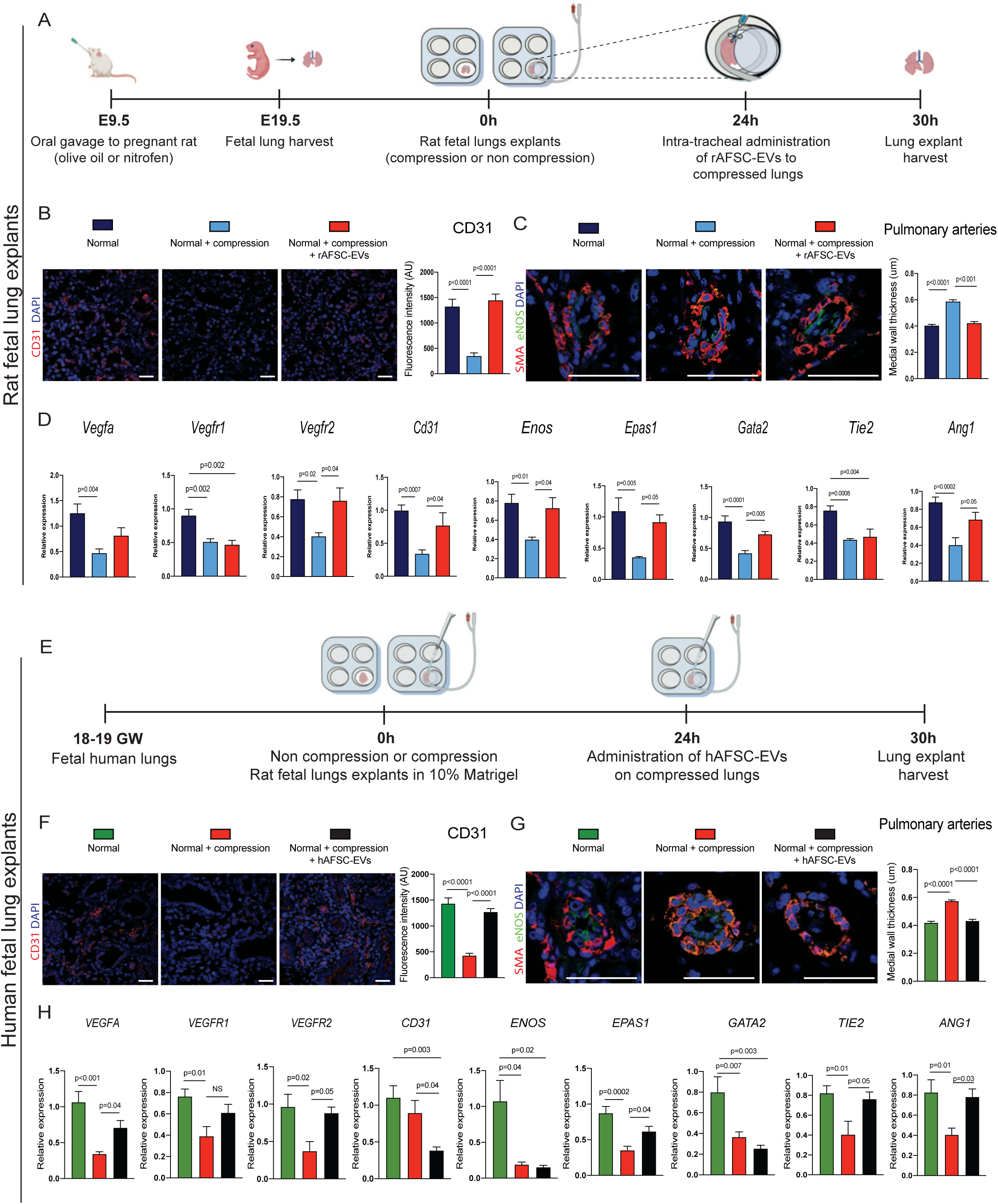
Administration of AFSC-EVs rescues lung vascular development in rat and human fetal lung explants. (A) Timeline for AFSC-EV *ex vivo* studies in rat fetal lung explants. (B) Representative images of immunofluorescence staining for CD31 in rat fetal lung explants (n = 5 in each group). Scale bar = 50 µm. Group comparison of CD31 fluorescence intensity (AU; 10 fields per biological replicate). (C) Representative immunofluorescence images of pulmonary arteries co-stained with SMA (red), eNOS (green), and DAPI (blue) in rat fetal lung explants (n = 5 in each group). Group comparison of medial wall thickness (10 fields per sample containing a minimum of 3 pulmonary arteries each). Scale bar = 50 µm. (D) Group comparison of gene expression of key angiogenic markers described in Fig. 1E relative to GAPDH for fetal rat lung explants (n ≥ 6 in each group). (E) Timeline for AFSC-EV *ex vivo* studies in fetal human lung explants. (F) Representative immunofluorescence images of CD31 in human fetal lung explants (n = 5 in each group). Group comparison for CD31 intensity (AU; 10 fields per biological replicate). Scale bar = 50 µm. (G) Representative images of pulmonary arteries co-stained for SMA (red), eNOS (green), and DAPI (blue) in human fetal lung explants (n = 5 in each group). Group comparison of medial wall thickness from 3-10 pulmonary arteries per field (µm). Scale bar = 50 µm. (H) Group comparison of gene expression of angiogenic markers in fetal human lung explants (n = 6 in each group). (B-H) Data are shown as mean ± SD. Groups were compared using one-way ANOVA (Tukey post-test) for (C; F = 53.68), (D; *Vegfa*, F = 6.50; *Vegfr1*, F = 9.92; *Vegfr2*, F = 5.40; *Cd31*, F = 9.59; *Enos*, F = 5.71; *Epas1*, F = 7.20; *Gata2*, F = 18.04; *Tie2*, 11.56; *Ang1*, F = 10.31), (G; F = 85.68), and (H; *VEGFA,* F = 15.71; *VEGFR2,* F = 5.10; *CD31,* F = 7.12; *EPAS1,* F = 13.11; *GATA2,* F = 8.25; *ANG1,* F = 6.29; and *TIE2,* F = 5.36); and Kruskal-Wallis (post hoc Dunn’s nonparametric comparison) for (B; U = 32.23), (F; U = 43.59), (H; *VEGFR1,* U = 8.03 and *ENOS,* U = 8.83) according to Shapiro-Wilk normality test. Only p-values < 0.05 are shown.

### Intra-amniotic administration of AFSC-EVs rescues vascular development in fetal rat lungs *in vivo*

Towards translation of this therapeutic approach, we investigated whether antenatal administration of AFSC-EVs to fetal lungs could rescue vascular development *in vivo*. To this end, we used our well established EV delivery method, whereby fetal rats receive an intra-amniotic (IA) injection of either saline or rAFSC-EVs at E18.5, as described ^21,25^ (**Figure 5A)**. Fetal rat lungs were harvested at E21.5, allocated into control+saline, CDH+saline and CDH+rAFSC-EV groups, and evaluated for number of airspaces. Only nitrofen exposed fetuses with diaphragmatic defect were selected for the CDH groups, as we did not find lung vascular abnormalities in nitrofen exposed fetuses without CDH (**Suppl.Fig. 3A-D**). We observed that in comparison to control+saline, CDH+saline fetal lungs had decreased number of airspaces that were rescued back to normal following intra-amniotic injection of rAFSC-EVs (**Suppl.Fig. 1E**). Moreover, we observed that in comparison to control+saline, CDH+saline fetal lungs had decreased vascular density and increased medial wall thickness of the pulmonary arteries (**Figure 5B-C**). These parameters were rescued back to control levels following rAFSC-EV IA administration (**Figure 5B-C**). Moreover, we found that compared to control+saline lungs, CDH+saline lungs had decreased expression of *Vegfa, Vegfr1, Vefgr2, Cd31, Enos, Epas1, Ang1* and *Tie2*. IA administration of rAFSC-EVs rescued the expression of all angiogenic markers back to normal levels in CDH fetal lungs (**Figure 5D**). Similarly, we observed that CDH lungs had decreased protein expression of VEGFR2, CD31, ENOS, EPAS1, GATA2, TIE2 and ANG1 and that IA injection of rAFSC-EVs restored the expression of VEGFR2, CD31, ENOS, EPAS1, GATA2 and TIE2 back to normal levels (**Figure 5E**). We also confirmed that compared to control+saline, lungs of CDH+saline fetuses had downregulated expression of markers of the Hippo signaling pathway that was rescued by the intra-amniotic administration of AFSC-EVs (**Figure 5F**). To explore the mechanism behind the beneficial effects of rAFSC-EVs in lungs of fetal rats with CDH, we interrogated miRwalk and identified miRNAs involved in biological processes that modulate vascular development and Hippo signaling pathway and that are known to be contained in the rAFSC-EV cargobased on previously established dataset (**Figure 6**) ^20^.

**Figure 5.**
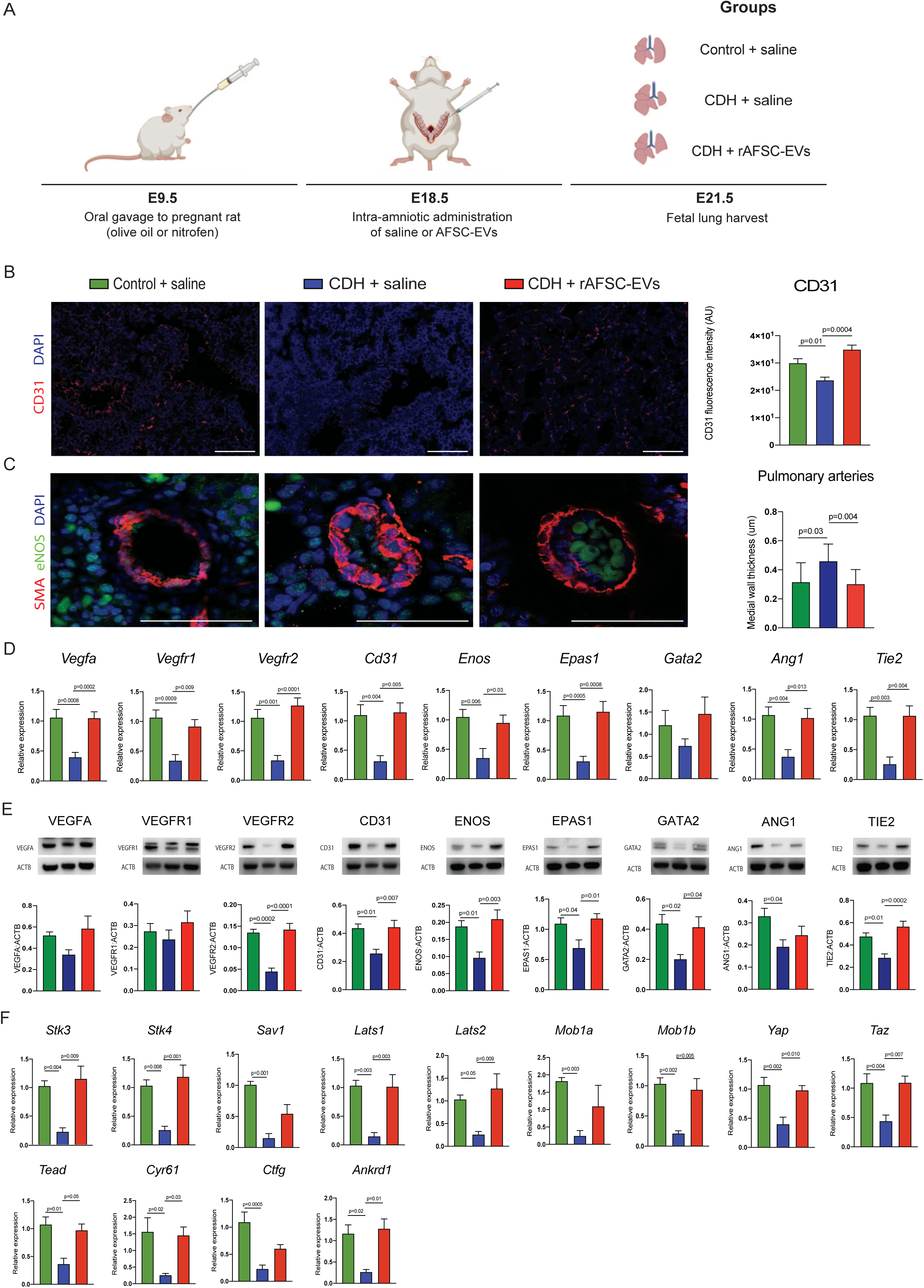
Antenatal administration of AFSC-EVs rescues lung vascular development and Hippo pathway signaling in an *in vivo* fetal rat CDH model. (A) Timeline of *in vivo* studies in fetal rats. (B) Representative immunofluorescence images of CD31 in rat fetal lungs (n ≥ 5 in each group). Scale bar = 50 µm. Group comparison of CD31 fluorescence intensity (AU; 10 fields per biological replicate). (C) Representative images of pulmonary arteries co-stained for SMA (red), eNOS (green), and DAPI (blue) in fetal rat lungs (n ≥ 5 in each group). Group comparison for medial wall thickness from 3-10 pulmonary arteries per field (µm). Scale bar = 50 µm. (D) Group comparison of gene expression for angiogenic markers relative to GAPDH (n ≥ 5 in each group) and (E) Representative Western blotting images and quantification of angiogenic factors relative to ACTB in fetal rat lungs (n ≥ 6 in each group). (F) Group comparison of gene expression of Hippo pathway factors relative to GAPDH fetal rat lungs (n = 6 in each group). (B-F) Data are shown as mean ± SD. Groups were compared using one-way ANOVA (Tukey post-test) for (B; F = 11.98), (C; F = 7.25), (D; *Vegfa*, F = 14.07; *Vegfr1*, F = 10.38; *Vegfr2*, F = 18.51; *Cd31*, F = 12.02; *Enos*, F = 7.32; *Epas1*, F = 14.80; *Gata2,* F = 1.32; *Ang1*, F = 8.14; and *Tie2,* F = 9.56), and (E; VEGFA, F = 2.32; VEGFR1, F = 0.77; VEGFR2, F = 21.75; CD31, F = 6.91; ENOS, F = 8.26; EPAS1, F = 5.58; GATA2, F = 5.06; ANG1, F = 3.49; and TIE2, F = 11.86) according to Shapiro-Wilk normality test. Only p-values < 0.05 are shown.

**Figure 6.**
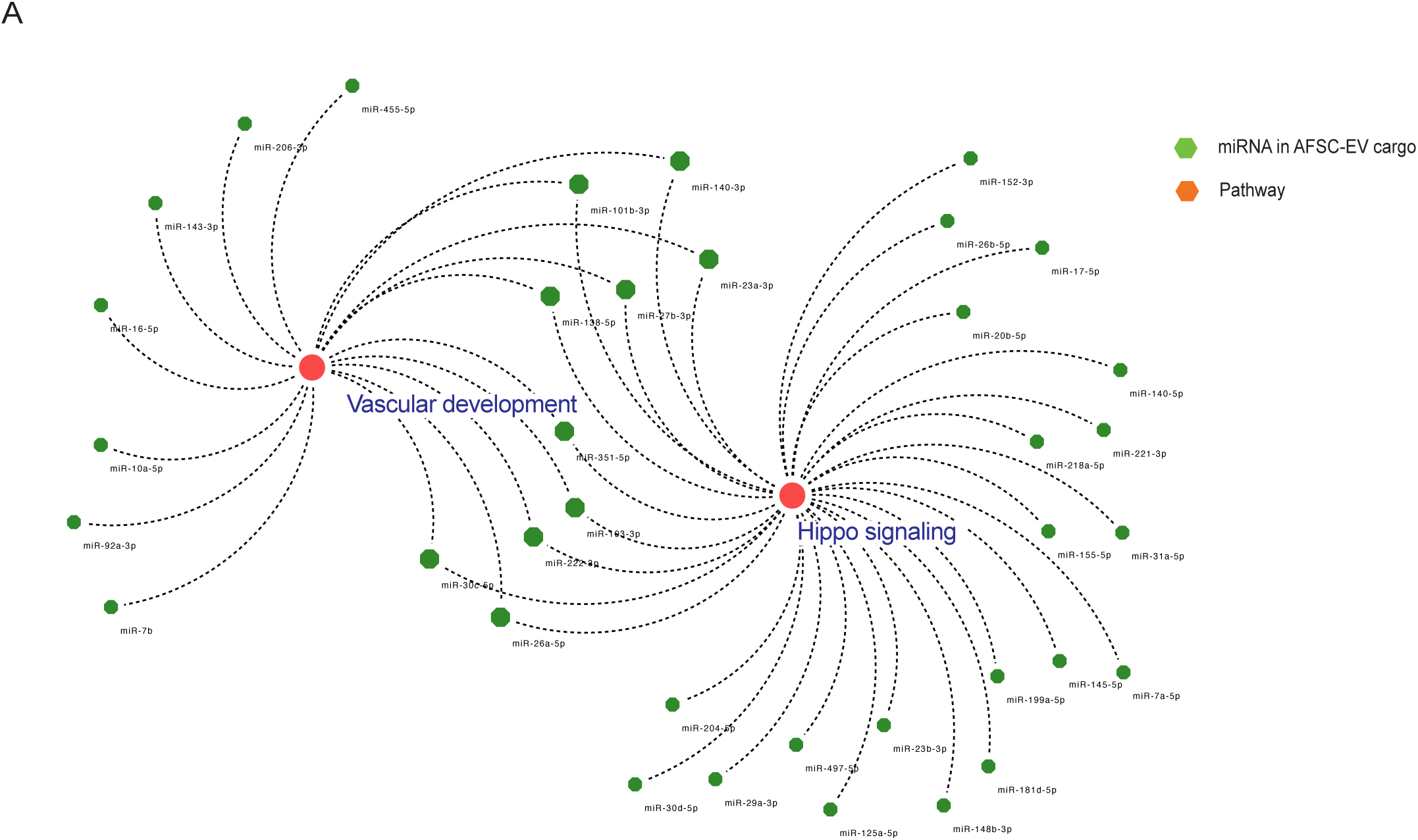
AFSC-EV cargo contains miRNAs that modulate vascular development and Hippo signaling pathway. (A) Network showing AFSC-EV miRNAS that independently or codependently regulate both angiogenesis and Hippo pathway biological processes.

## Discussion

Compression of fetal lungs is a primary characteristic of babies with CDH and a known factor for the pathogenesis of the impaired fetal lung development in these babies. This concept has recently been confirmed by studies that used mechanical compression systems and proved that compressed fetal lung explants and lung organoids have impaired branching morphogenesis and decreased epithelial cell differentiation^14-16^. Compared to *in vivo* animal models of CDH, these systems had the ability to induce controlled compression even in healthy fetal lungs and study the effects of compression in isolation. Our study builds on these previous studies and reveals that mechanical compression plays an essential role also in the disruption of fetal lung vascular development. Specifically, our findings showed that mechanical compression alone is essential for vascular remodeling in nitrofen-exposed lungs, as well as in normal healthy lungs. Compared to the other models, our novel static micro-compression system considered all the forces that play a role in a fetus with CDH. The calculations were based on the fetal lung pulmonary fluid volume, density, viscosity and pressure; the fetal breathing movements, both clustered and not; the tracheal length and diameter; and the weight of the pup. This model enabled us to compress the left lung at a translationally relevant time point of lung development and reproduce similar pulmonary hypoplasia features of the ones observed in human babies with left-sided CDH. Using this system, we could study the effects of nitrofen in isolation. In fact, previous studies using the *in vivo* nitrofen model proved that rat fetuses with CDH had features of lung vascular remodeling, such as hypertrophy of the medial wall thickness in pulmonary arteries ^28-30^. However, these studies did not show the isolated effects of mechanical compression, given that in the nitrofen model of CDH all fetuses are inevitably exposed to nitrofen. Using our novel *ex vivo* model, we overcome this limitation by compressing normal fetal lungs and comparing them to nitrofen exposed fetal lungs. Our experimental findings revealing that mechanical compression induces fetal lung vascular remodeling are supported by clinical evidence. On the one hand, babies with larger diaphragmatic defect and liver up position have more severe pulmonary hypertension and increased need for extracorporeal life support (ECLS) than those with smaller defects ^31,32^. This suggests that the degree of mechanical compression exerted by the herniated organs is directly proportional to the severity of pulmonary hypoplasia and predicts pulmonary hypertension. On the other hand, babies with CDH and a hernia sac, i.e. a thin membrane that contains the herniated organs, are known to have less severe pulmonary hypertension and lower ECLS requirement compared to babies with CDH without a sac ^33^. These data indicate that an attenuated mechanical compression to the lungs of fetuses with CDH is associated with less severe lung vascular remodeling and postnatal pulmonary hypertension.

Our study also revealed that mechanical compression to fetal lungs is associated with a dysregulation of the Hippo signaling pathway. This signaling pathway is a conserved mechanosensitive pathway that regulates cell proliferation through response to mechanical stimuli and is critical for organ development ^34-40^. The Hippo signaling pathway has been reported to be dysregulated in rodent models of CDH ^39, 41-43^. Serapiglia et al. reported that fetal mouse with conditional depletion of Yap from the lung epithelium had reduced lung basal cell population in comparison to fetal lungs from wild-type mouse ^42^. Kahnamoui et al. demonstrated that the Hippo signaling effector protein Yap is inactive in nitrofen-induced hypoplastic lungs in comparison to control lungs ^43^. The downregulation of upstream and downstream effectors of the Hippo signaling pathway in our *ex vivo* model was exclusively dependent on compression, concomitant to the impaired vascular development in compressed rat and human fetal lungs. Studies have shown that Yap modulates endothelial cell sprouting through several key ligands/receptors such as Vegf/Vegfr and angiopoietin/Tie2 and it is required for regenerative lung growth after injury ^37,44^. Therefore, it is reasonable to state that mechanical compression triggers a cascade of biological events concurrent with the disruption of the Hippo signaling pathway leading to aberrant lung vascular development.

The multifactorial pathogenesis of arrested lung development in CDH necessitates an antenatal therapy that addresses all aspects of impaired fetal lung growth, maturation, and vascularization ^8^. We have previously shown that AFSC-EVs restore fetal lung branching morphogenesis and epithelial cell differentiation in rodents and human models of CDH ^20-22^. The current study provides evidence that AFSC-EV administration to rat and human compressed fetal lungs rescues also fetal lung vascular development. Other groups have shown similar effects in experimental CDH models upon administration of EVs derived from mesenchymal stromal cells (MSC-EVs) ^45,46^. Specifically, MSC-EV administration was reported to restore endothelial cell viability, improve pulmonary arterial contractility, and mitigate extracellular matrix remodeling in models of CDH ^45,46^. There is also evidence that an EV-based therapy improves lung vascular development and prevents pulmonary hypertension in experimental models of bronchopulmonary dysplasia, a neonatal lung condition that affects premature babies and is similar to pulmonary hypoplasia secondary to CDH ^47-51^. In these studies, EV-based therapies were reported to improve lung vascular density, attenuate vascular remodeling, and prevent pulmonary hypertension ^47-51^. This experience with EVs in a neonatal condition mirrors the beneficial effects demonstrated by EV-based therapy in adult preclinical models of pulmonary arterial hypertension ^52,53^. The regenerative effects of EV administration in these studies have been ascribed in part to the release of miRNAs contained in the EV cargo ^53-56^. This is in line with our previous mechanistic studies, where we demonstrated that one potential mechanism of action of AFSC-EVs is through the delivery of miRNAs to the fetal lungs ^20,21^. In the current study, we confirmed that the AFSC-EV cargo is enriched with some miRNAs that are involved in vascular development and Hippo signaling pathway. This is relevant as miRNAs are critical mediators of fetal lung vascular development ^57^ and their expression has been reported to be dysregulated in human newborns with CDH with pulmonary hypertension ^58,59^.

Our results are important from a translational standpoint as they prove that EV administration is beneficial even in compressed lungs. Mechanical compression in CDH is relieved only upon surgical repair of the diaphragmatic defect, which occurs only postnatally. At present, the only antenatal therapy that can be offered to selected fetuses with CDH is FETO, whereby a balloon is endoscopically deployed in the fetal trachea to cause distention of the hypoplastic lungs ^8^. A randomized controlled trial has shown that FETO improves survival in fetuses with CDH and severe pulmonary hypoplasia ^18,19^. However, there is evidence that FETO does not attenuate vascular remodeling and does not prevent pulmonary hypertension in these babies ^17,60^. For this reason, there have been various attempts to use pharmacological adjuncts that could be administered to the pregnant mother, reach the fetus transplacentally, and rescue normal vascular development. To this end, Sildenafil, a selective inhibitor of phosphodiesterase type 5 with vasodilatory effects on the pulmonary circulation, was trialled in fetuses with fetal growth restriction but failed the interim analysis due to high fetal demise ^61, 62^. An alternative that is now being tested in preclinical models of CDH is Treprostinil, a synthetic prostacyclin analog that crosses the placental barrier and has an anti-remodeling effect on the pulmonary vasculature ^63^. An EV-based therapy could be an optimal adjunct for FETO, as it can be delivered topically to the fetal lungs at the time of the balloon deployment and would address all aspects of fetal lung underdevelopment, including vascularization. We have preclinical evidence that AFSC-EVs can be delivered at the time of tracheal occlusion both *in vivo* in fetal rabbits ^20^ and *ex vivo* in fetal rat explants, as herein shown.

We acknowledge that this study has some limitations. Our ex vivo model of mechanical compression, like the other ex vivo models recently reported ^14-16^, does not fully replicate the dynamic environment of a human CDH fetal lung *in utero*. Nonetheless, our model allowed application of controlled pressure on both normal and nitrofen-exposed fetal lungs. Moreover, our data on the Hippo pathway are extrapolated from the compression model of healthy human fetal lungs and are not derived from lungs of fetuses with CDH. However, fetal lung specimens from babies with CDH are not available for several reasons, including the low rate of terminations for CDH and the prevalence of severe genetic abnormalities that do not represent the typical pulmonary hypoplasia phenotype. Furthermore, the promising effects exerted by AFSC-EVs on the fetal lung vasculature are based on morphometric changes in preclinical models. This limitation is due to challenges to assess pulmonary hypertension postnatally, as the available models are non-surviving. Lastly, the AFSC-EV effects on the lung vasculature could also be influenced by other bioactive molecules beyond miRNAs. Other small RNA species are in fact contained in the AFSC-EV cargo but at present their biological role remains unknown.

## Material and Methods

### Rat experiments

Following ethical approval from the Animal Care Committee at the Hospital for Sick Children (AUP#65210), pregnant Sprague-Dawley rats received 100mg of nitrofen (2,4-Dichlorophenyl 4-nitrophenyl ether; Sigma CAS#1836-75-5) in 1mL of olive oil at E9.5 to induce fetal pulmonary hypoplasia. In this model 100% of the litter develops pulmonary hypoplasia and 40-60% develop the diaphragmatic defect (CDH+). Pregnant rats assigned to the control group received olive oil (alone) at E9.5.

#### *Ex vivo* rat model

At E19.5, fetuses were harvested, CDH was confirmed, and only lungs from fetuses without CDH were selected. Control fetuses were harvested at the same timepoint. and allocated into the following groups (n=8-12): 1) Normal, 2) Normal + compression, 3) Nitrofen exposed; 4) Nitrofen + compression. Fetal rat lungs were cultured as explants in 10% Matrigel and subjected to either mechanical compression using our micro bioengineered system for 24 hours (described below) at 37°C or left non-compressed.

### Human experiments

#### *Ex vivo* human model

Fetal human lungs were obtained from terminations of healthy fetuses (n=7) with no CDH at 18-19 weeks of gestation following ethical approval from Mount Sinai Hospital Biobank, Toronto (REB#10-0128-E). Lung pieces weighing 1-1.7g were cultured as explants in 10% Matrigel, and randomly assigned to receive mechanical compression or left non-compressed.

#### Autopsy samples

Lung autopsies were obtained from fetuses with CDH (n=4) and sex- and age-matched controls (n=4) following ethical approval from The Hospital for Sick Children, Toronto, ON, Canada (REB#1000074888).

### Mechanical compression system

We established a novel bioengineered static micro-compression system that simulates the herniation of intraabdominal organs into the thoracic cavity as observed in CDH fetuses. Lungs from either rat human fetuses at the canalicular stage were chosen to perform mechanical compression, as this corresponds to the time-point of CDH diagnosis in human babies. We calculated the ideal load to apply on fetal lung explants by considering the various forces present in the uterine microenvironment such as the pulmonary fluid volume, density, viscosity and pressure; the fetal breathing movements, both clustered and not; the tracheal length and diameter; and the weight of the pup. The final equation was: ΔP = P* - PCDH, where P* is lung pressure at E19.5 and PCDH is the pressure exerted by the herniated abdominal organs. P* was calculated by taking in consideration the force exerted by fetal pulmonary fluid secretion (0.00152 Newton), upper airway resistance (0.2 cm H2O/mL/s), and fetal breathing movements. Thus, our mechanical compression system consists of applying a load of 0.22 kPa via the inflation of a 6 French Foley catheter balloon (RÜSCH Brillant, Teleflex medical) with 0.9mL buffered saline towards fetal lungs cultured in 10% Matrigel matrix (Corning^TM^) + culture medium (Gibco^TM^, DMEM) for 24h at 37°C.

#### *In vivo* rat model

At E18.5, pregnant rats were anesthetized, a vertical midline laparotomy was performed to expose uterine horns, 100 ul of either saline or AFSC-EVs was injected into the amniotic sac using a 30G needle, the fetuses were placed back into the abdominal cavity, and abdominal wall was closed. At E21.5, fetal lungs were harvested and allocated into the following groups (n=6-10): 1) control (healthy fetuses), 2) Nitrofen exposed fetuses without diaphragmatic defect (Nitrofen) and 3) CDH, nitrofen exposed fetuses with diaphragmatic defect.

### AFSC-EV isolation and characterization

For rat studies, EVs were derived from c-kit+ rat and human AFSC conditioned medium, and cultured in vitro until 80% confluence as described ^20,21^. Rat and human AFSC-EVs were isolated by ultracentrifugation and characterized by size, morphology and expression of canonical EV protein markers according to the International Society for Extracellular Vesicles as described^20,21,64^.

### Radial airspace count

Fetal rat and human lung samples were fixed in paraformaldehyde, paraffin-embedded, sectioned to 4 μm sections, and stained with hematoxylin and eosin staining (H&E). H&E stained lung sections were assessed for the number of airspaces (radial airspace count, RAC) ^65^. RAC analysis was performed on 20X scanned H&E fetal lung samples processed in CaseViewer Software and quantified in the whole lung section, as previously described ^20,21^.

### Immunofluorescence

Fixed lung sections from rat and human fetal lungs were incubated with primary antibodies for vascular density analysis (endothelial cells, CD31) and for medial wall thickness assessment of the pulmonary arteries (smooth muscle cell, SMA; and endothelial nitric oxide synthase, eNOS). Primary and secondary antibodies concentration are listed in Supplementary Figure 4. Leica SP8 lightning confocal microscope (Wetzlar, Germany) was used to image samples using similar laser power and exposure between conditions. Vascular density was quantified by assessing CD31 fluorescence intensity on fetal lungs with HistoQuant in QuantCenter Imaging Software (3D Histech) and covered >100 50×50 μm fields.

### Medial wall thickness

To identify the muscle layer of pre-acinar arteries in fetal lung sections, we co-stained fetal lungs with factors for smooth muscle cells (smooth muscle actin, SMA) and endothelial cells (endothelial nitric oxide, eNOS). For each lung section, 10 to 30 pulmonary arteries were identified and measured using HistoQuant in QuantCenter Imaging Software (3D Histech) at 400x magnification. Medial wall thickness (MWT) was assessed by the following equation: External diameter – Internal diameter / External diameter (ED – ID / ED), as described ^66, 67^.

### Gene expression analysis

Fetal rat and human lung explants were harvested and frozen at -20°C until analysis. Total RNA was isolated using Trizol reagent following supplier recommended protocols (ThermoFisher Scientific, Waltham, MA). Purified RNA was quantified using a NanoDrop™ spectrophotometer (ThermoFisher Scientific, Waltham, MA) and cDNA synthesis was performed with 500ng quantified RNA (superscript VILO cDNA synthesis kit, ThermoFisher Scientific, Waltham, MA). qPCR experiments were conducted with SYBR™ Green Master Mix (Wisent, Saint-Jean-Baptiste, QC) for 45 cycles (denaturation: 95°C, annealing: 58°C, extension: 72°C) using the primer sequences designed for rat and human lungs reported in table S2 and S3, respectively. Melt curve plots were used to determine target specificity of the primers. ΔΔCT method was used to determine normalized relative gene expression.

### Protein expression analysis

Protein from rat and human fetal lung explants was isolated by incubation with cell extraction buffer (ThermoFisher Scientific, Waltham, MA) supplemented with protease inhibitors (Sigma Aldrich, Missouri, MO), and sonicating for 3 cycles of 10 seconds each. Protein was quantified using the Pierce BCA Protein Assay (ThermoFisher Scientific, Waltham, MA), and 15μg of protein from each sample was processed and probed for the vascular markers. B-Actin was used as a loading control.

### Statistical analysis

Groups were compared using two-tailed Student t-test, Mann-Whitney test, one-way ANOVA (Tukey post-test), or Kruskal-Wallis (post-hoc Dunn’s non-parametric comparison) test according to Gaussian distribution assessed by D’Agostino Pearson omnibus normality test. P value <0.05 was considered significant. All statistical analyses were produced using GraphPad Prism^®^ software version 8.4. Differential gene expression analysis was conducted using BioConducter R (3.15) package MAST (1.22.0) between two conditions, with adjusted p-value <0.05 and log2(fold change) > |0.5| considered as significant for snRNA-seq experiments.

## Acknowledgements

The authors would like to thank A. Ijaz and C. Yu, for their help with the miRNA network analysis. We are indebted to P. De Coppi for providing rat AFSCs in kind, the Lab Animal Service core facility at the Hospital for Sick Children, Nanoscale Biomedical Imaging Facility and the Imaging Facility at the Hospital for Sick Children, Toronto for their services. We would also like to show gratitude the Mount Sinai Hospital Biobank, Toronto.

Funding: Canadian Institutes of Health Research (CIHR)-CIHR Project Grant (175300, AZ; 497959, AZ), American Pediatric Surgical Association Grosfeld Scholarship (AZ), SickKids Congenital Diaphragmatic Hernia Fund (R00DH00000, AZ), American Thoracic Society (RP-2020-26, RLF), CIHR Fellowship (187855, RLF; 176535, KK), and German Research Foundation DO (466815475, FD).

## Author contributions

M.S.G., L.A., and A.Z. conceived and established the *ex vivo* model, R.L.F., L.A. and A.Z. conceived and designed the *in vivo* studies. R.L.F., K.K., L.A., F.D., M.O. and S.G. conducted the experiments. A.Z. supervised the experiments. R.L.F., K.K. and L.A. analyzed the data. R.L.F., L.A. and A.Z. wrote the manuscript.

## Data Availability

All data generated and analyzed during the present study are included in this published article and its supplementary information files.

