## Supplementary data for "Fetal lung vascular development is disrupted by mechanical compression and rescued by administration of amniotic fluid stem cell extracellular vesicles via regulation of the Hippo signaling pathway"

### Supplementary Figure 1

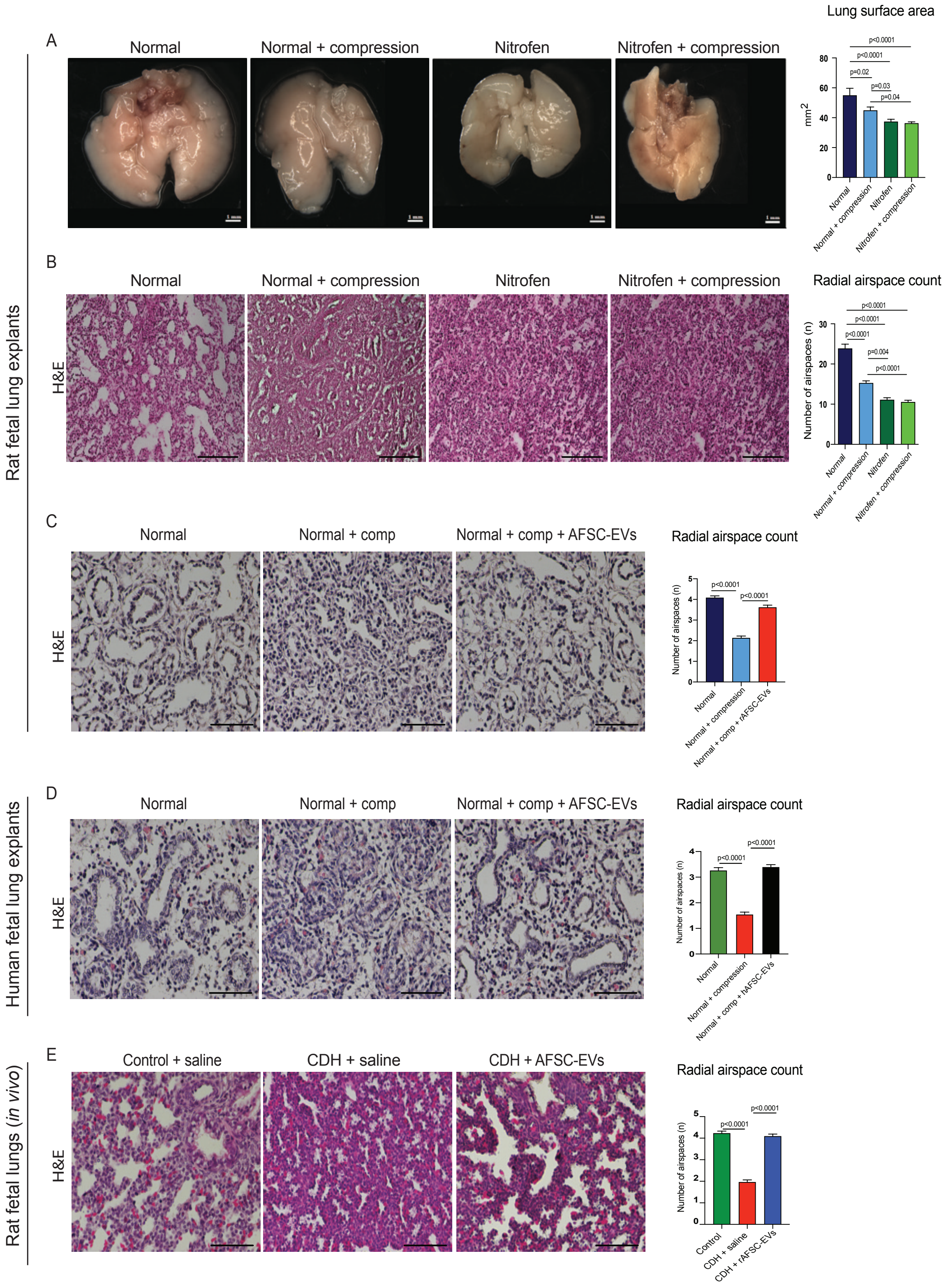

**Supplementary figure 1. Mechanical compression affects branching morphogenesis in fetal rat and human lungs.** (A) Representative brightfield microscopy images of fetal rat lung explants. Group comparison of lung surface area ($mm^2$; ImageJ software; n=5 biological replicates per condition;). Scale bar = 1mm. (B) Representative histology images (hematoxylin/eosin staining) of fetal rat lung explants with group comparison of RAC (n = 5 in each group). Scale bar = 50 µm. (C) Representative histology images (hematoxylin/eosin staining) of fetal rat lung explants and group comparison of RAC (n=5 in each group). Scale bar = 50 µm. (D) Representative histology images (hematoxylin/eosin) of fetal human lung explants and group comparison of RAC (n = 5 in each group). Scale bar = 50 µm. (E) Representative histology images (hematoxylin/eosin) of fetal rat lungs at E21.5 with group comparison of RAC (n=5 in each group). Five fields per sample. Scale bar = 50 µm. (A-E) Data are shown as mean ± SD. Groups were compared using one-way ANOVA (Tukey post- test) for (A); and Kruskal-Wallis (post hoc Dunn's nonparametric comparison) for (B; F = 14.24), (C; H = 109.4), (D; H = 109.4) and (E; H = 124.2) according to Shapiro-Wilk normality test. Only p-values < 0.05 are shown.

Supplementary Figure 2

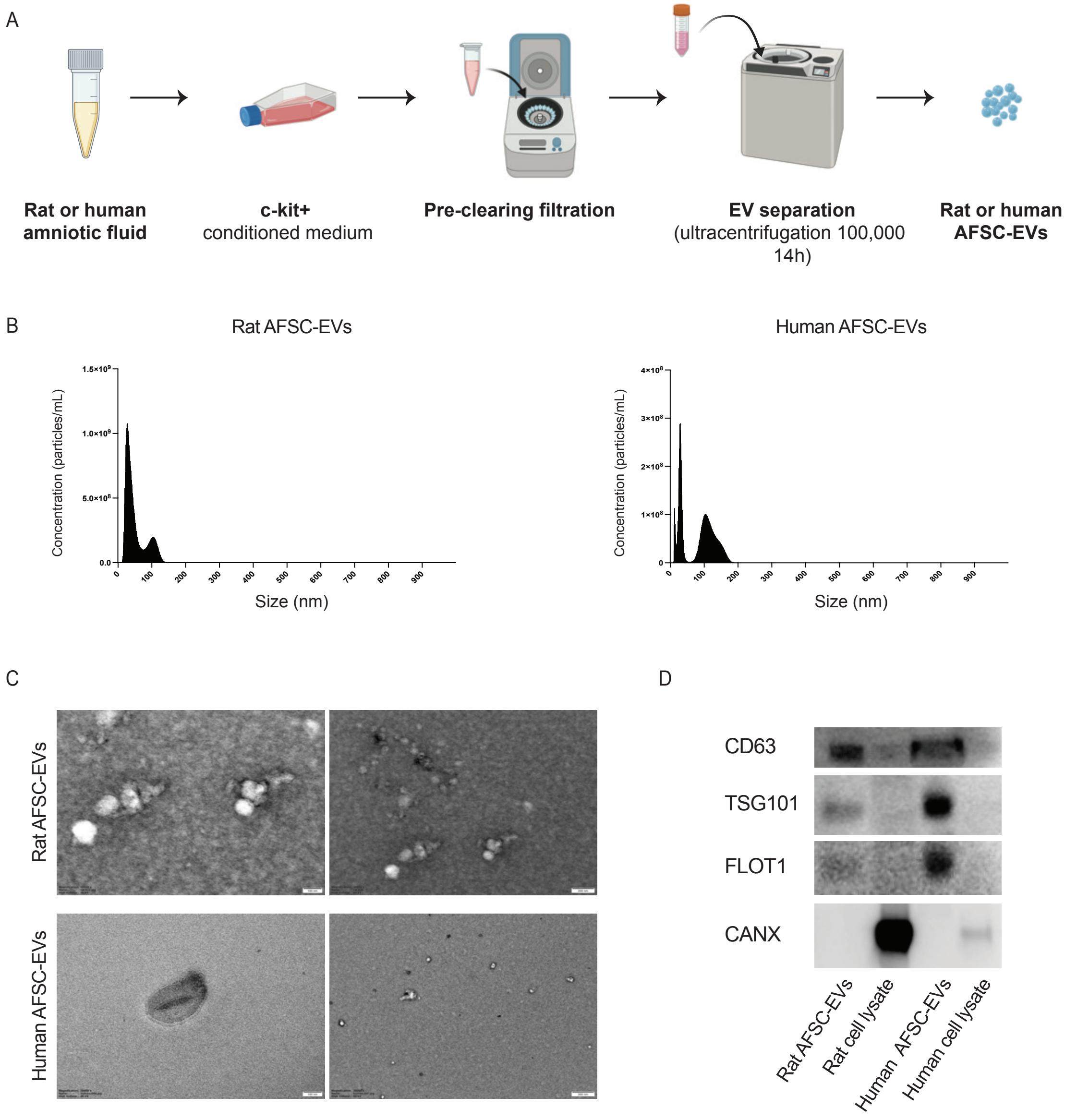


**Supplementary figure 2. Rat and human AFSC-EV isolation and characterization.** (A) Rat or human AFSC-EV isolation by ultracentrifugation (B) Representative plot of the average size distribution of rat and human AFSC-EVs visualized using nanoparticle tracking analysis. Data are representative of three 40-second videos of each EV preparation. X-axis = size distribution (nm), y-axis = concentration (particles/mL). (C) Representative transmission electron microscopy photos of rat and human AFSC-EVs. Scale bars: 100 nm for 5000x images and 200nm for 3000X images.

(C) Canonical EV markers CD63, TSG101 and Flot1 were positively detected in rat and human AFSC-EVs and were negative for the endoplasmic reticulum marker, Calnexin.

Supplementary Figure 3

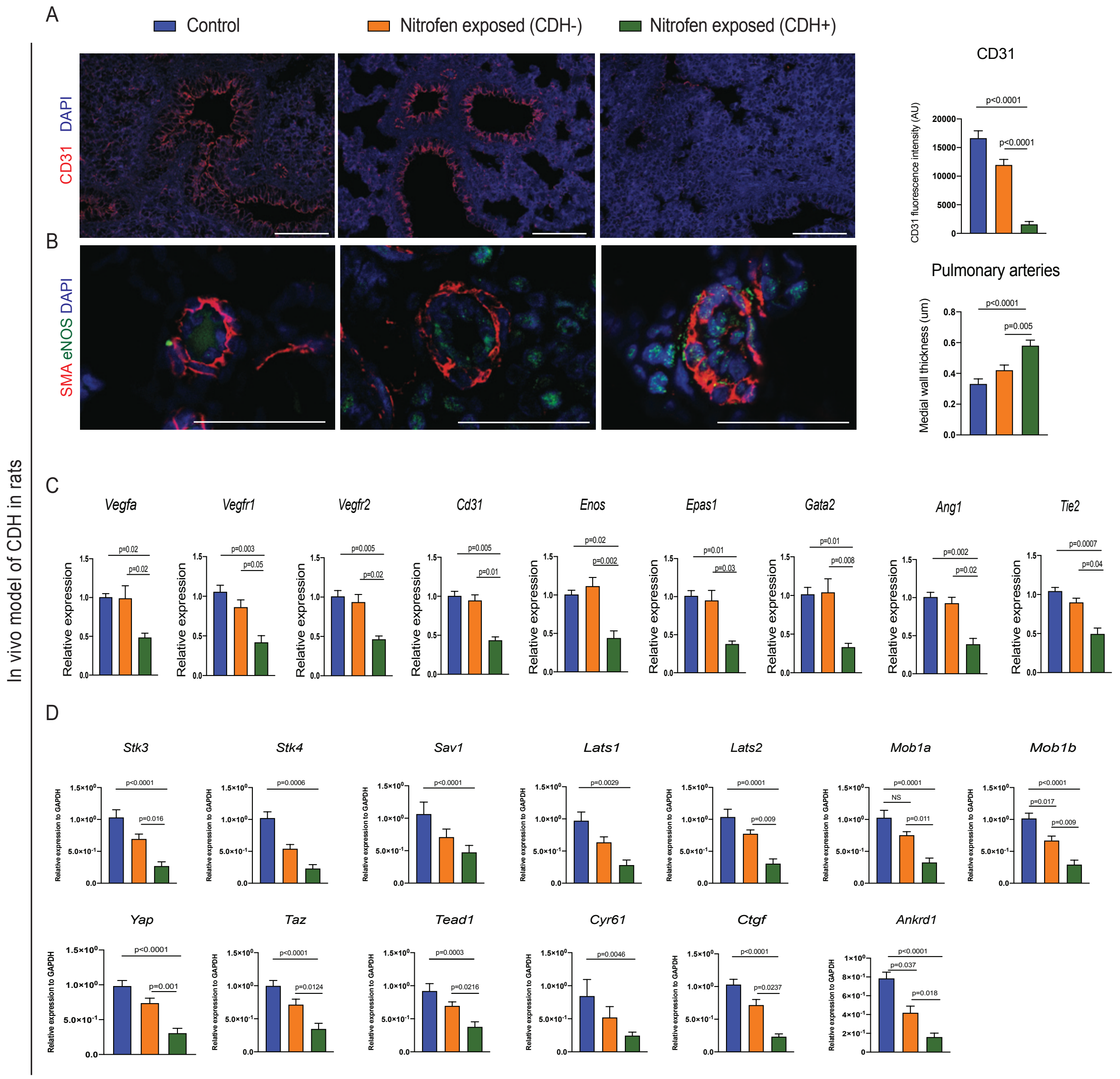


**Supplementary figure 3. Nitrofen exposure alone (without CDH) does not impair vascular development and Hippo pathway gene expression in the *in vivo* rat model.** (A) Representative immunofluorescence images of CD31 in rat fetal lungs at E21.5 (n = 5 in each group). Group comparison of CD31 fluorescence intensity (AU; 10 fields per biological replicate). Scale bar =

50 μm. (B) Representative images of pulmonary arteries co-stained for SMA (red), eNOS), and DAPI (blue) in fetal rat lungs (n = 5 in each group). Group comparison of medial wall thickness from 3-10 pulmonary arteries per field (μm). Scale bar = 50 μm. (C) Group comparison of gene expression of key angiogenic markers relative to GAPDH in fetal rat lungs (n = 5 in each group). (D) Group comparison of gene expression of all upstream and downstream Hippo signaling pathway factors in relative to GAPDH fetal rat lungs (n = 5 in each group). (A-D) Data are shown as mean ± SD. Groups were compared using one-way ANOVA (Tukey post- test) for (B; F = 14.58), (C; *Vegfa*, F = 8.60; *Vgefr1*, F = 15.72; *Vegfr2,* F = 16.54; *Cd31,* F = 29.28; *Enos*, F = 16.82; *Epas*, F = 15.93; *Gata2*, F = 11.38; *Tie2*, F = 18.51; *Ang1*, F = 23.82), and (D; *Stk3,* F = 18.21; *Lats1,* F = 10.88, *Lats2,* F = 16.91; *Mob1a,* F = 17.30; *Mob1b,* F = 24.55*; Yap,* F = 20.71*; Taz,* F = 16.02*; Tead,* F = 10.99; and *Ankdr1,* F = 26.53); and Kruskal-Wallis (post hoc Dunn's nonparametric comparison) for (A; H = 71.45) and (D; *Stk4, H = 13.82; Sav1,* F = 8.17*; Cyr61*, F = 10.03; and *Ctfg,* F = 20.86) according to Shapiro-Wilk normality test. Only p-values < 0.05 are shown.

**Table 1. Antibodies utilized for immunofluorescence and Western blotting.**

| Gene name | Antibody | Antibody host | Manufacturer/Catalogue # | Application | Dilution Factor |
|---|---|---|---|---|---|
| Vascular endothelial growth factor | VEGF | Mouse | Santa-Cruz/Sc7269 | WB | 1:500 |
| Platelet endothelial cell adhesion molecule | CD31/PECAM-1 | Goat | R&D/AF3628 | WB, IF | 1:250 (WB), 1:25 (IF) |
| Vascular endothelial growth factor receptor 1 | VEGFR1 | Rabbit | Abcam/ab32152 | WB | 1:250 |
| Vascular endothelial growth factor receptor 2 | VEGFR2 | Rabbit | Abcam/ab39638 | WB | 1:500 |
| Endothelial PAS domain-containing protein 1 | EPAS1 | Rabbit | Novus/NB100-122 | WB | 1:250 |
| Endothelial nitric oxide synthase | ENOS | Mouse | Abcam/ab76198 | WB, IF | 1:1000 (WB), 1:100 (IF) |
| Endothelial transcription factor GATA-2 | GATA2 | Rabbit | Abcam/ab109241 | WB | 1:250 |
| Angiopoietin-1 | ANG1 | Rabbit | Novus/NBP1-90169 | WB | 1:500 |
| Angiopoietin-1 receptor | TIE2 | Rabbit | Thermo Fisher/19157-1-AP | WB | 1:500 |
| Actin alpha 2/alpha smooth muscle actin | ACTA2/A-SMA | Rabbit | Abcam/ab124964 | IF | 1:200 |
| CD63 | CD63 | Mouse | Abcam/ab59479 | WB | 1:500 |
| Tumor susceptibility 101 | TSG101 | Mouse | Santa-Cruz/Sc79644 | WB | 1:1000 |
| Flotillin-1 | Flotillin-1 | Mouse | BD/610820 | WB | 1:500 |
| Calnexin | Calnexin | Rabbit | Abcam/ab22595 | WB | 1:1000 |
| Not applicable | IgG Alexafluor-488 | Rabbit | CST/4412S | IF | 1:1000 |
| | IgG Alexafluor-555 | Mouse | CST/4409 | IF | 1:1000 |
| | IgG HRP | Mouse | CST/7076S | WB | 1:1000 |
| | IgG HRP | Rabbit | CST/7074 | WB | 1:1000 |

**Table 2. Primer sequences used in human experiments.**

| Target | Forward primer | Reverse primer |
|---|---|---|
| *STK3* | 5'- CCTTTCCCGCCCTTAAGTAT-3' | 5'- CCAAATCTTTTCAAGAGACCGTAGT -3' |
| *STK4* | 5'-TGCCAGATGCCAGAAATTTATAC-3' | 5'- ATGTTGCTCAGGCTGGTATC-3' |
| *SAV 1* | 5'- GAGAGAGAGAGAGCCAGACCTT-3' | 5'- TCACGCACCTTCCCAATAC-3' |
| *LATS1* | 5'- CAGCAGCTGCCAGACCTATTA -3' | 5'- TGAGCTTGAACAAATGCTGATATAC -3' |
| *LATS2* | 5'- CTGGTGGGGACTCCAAACTAC-3' | 5'- GCTGGGTTTCTGTGGGAGTA-3' |
| *MOB1A* | 5'- GGTGGGAGCAACCAAAGTTA -3' | 5'- CCTTTTCTCAGCCAAAAGCTACTA -3' |
| *MOB1B* | 5'- CCAGAGGGGTCTCACCAGTAT -3' | 5'- CCGGACAGCTCTCCTCTGTA -3' |
| *YAP1* | 5'- CAGAACCGTTTCCCAGACTAC-3' | 5'- GTCTGCCTGAGGGCTCTATAA-3' |
| *TAZ* | 5'- CAATTTCCATGGGGGACTTA-3' | 5'- CTTGTTTTGCTTTTGGGGTACTA-3' |
| *TEAD1* | 5'- CTCTTCCCTGCCAGCACTATAC-3' | 5'- GGTGCATGACTTCAGTGAAACTAC-3' |
| *CYR61* | 5'- GGGCAGACCCTGTGAATATAA -3' | 5'- TCCATGGGGTCCTTGATACTA -3' |
| *CTGF* | 5'- TGACATCTTTGAATCGCTGTACTA -3' | 5'- CCAGTGTCTGGGGTTGATAGA -3' |
| *ANKRD1* | 5'- AAAAATTAGCGCCCGAGATA -3' | 5'- TCAGGAGTCGGATCATCTTATAG -3' |
| *VEGFA* | 5'- TGAGGAGTCCAACATCACCA -3' | 5'- TTTCTTGCGCTTTCGTTTTT -3' |
| *VEGFR1* | 5'- ACAACTCGGTGGTCCTGTACT -3' | 5'- CTGGCTCCCATGGAAAGATA -3' |
| *VEGFR2* | 5'- AAACGCTGACATGTACGGTCTA -3' | 5'- CGCTTGGATAACAAGGGTACT -3' |
| *CD31* | 5'- ATGGCAACAAGGCTGTGTACT -3' | 5'- ATGCTGCTGACCTTGGATATG -3' |
| *ENOS* | 5'- CAGGTGGGATGCGAACTTA -3' | GAGCTGTGCCTCCGTTTATAG -3' |
| *EPAS1* | 5'- TGCACCAAGGGTCAGGTAGT -3' | 5'- CAGGTTGCGAGGGTTGTAGA -3' |
| *GATA2* | 5'- TGAAGATGGAAAGTGGCAGTC -3' | 5'- GGAAGAGTCCGCTGCTGTAGT -3' |
| *TIE2* | 5'- TCCCGAGGTCAAGAGGTGTA -3' | 5'- CACAAGTCATCCCGCAGTAG -3' |
| *ANG1* | 5'- GAAGGGAACCGAGCCTATTC -3' | 5'- AGGGCACATTTGCACATACA -3' |

**Table 3. Primer sequences used in rat experiments.**

| Target | Forward primer | Reverse primer |
|---|---|---|
| *Stk3* | 5’- CGATGGCAAAACGCAATACT-3' | 5’- TGTTGGTGGTGGGTTTGTAG-3’ |
| *Stk4* | 5’- GCCCACATGTCGTCAAATACTA-3' | 5’- CGCCTTGATATCTCGGTGTAT -3' |
| *Sav1* | 5'- AAAGGAAGCGGACGAGAG-3' | 5'- TAGGGAGGAGGTGGGAGTAGA -3' |
| *Lats1* | 5'- CAGCAGCTGCCAGACCTATTA -3' | 5'- TGAGCTTGAACAAATGCTGATATAC -3' |
| *Lats2* | 5'- CTGGTGGGGACTCCAAACTAC-3' | 5'- GCTGGGTTTCTGTGGGAGTA-3' |
| *Mob1a* | 5'- GGTGGGAGCAACCAAAGTTA -3' | 5'- CCTTTTCTCAGCCAAAAGCTACTA -3' |
| *Mob1b* | 5'- CCAGAGGGGTCTCACCAGTAT -3' | 5'- CCGGACAGCTCTCCTCTGTA -3' |
| *Yap1* | 5’- GCATGGTGTGCCTGGTTATAC-3' | 5’- ATCTGCATGCTACCCACTACAG-3’ |
| *Taz* | 5’- GCCTGCCATGAACACAGATA-3' | 5’- TCCATGTTGCTGAGGAAGTCT-3’ |
| *Tead1* | 5'- AGAATGGCCGATTCGTGTA-3' | 5'- GCTGTGCTCCATGCTCACTAT-3' |
| *Cyr61* | 5'- AAGAAATACCGGCCCAAATAC -3' | 5'- GGATGCGGGCAGTTGTAGT -3' |
| *Ctgf* | 5'- TGACAGGGGAGGGACATTATAG -3' | 5'- TGCCACAAGCTGTCCAGTCTA -3' |
| *Ankrd1* | 5'- GAACCGGAGCCTGAAATTATTA -3' | 5'- GCTGTGGATTCCAGCATATC -3' |
| *Vegfa* | 5'- CAGGAGTACCCCGATGAGATAG -3' | 5'- GATCCGCATGATCTGCATAGT -3' |
| *Vegfr1* | 5'- GAGGACGCAGGGGACTATAC -3' | 5'- AACCACACGGGCCTCTACT-3' |
| *Vegfr2* | 5'- TGTCCCAGGGCTGACTCTAC -3' | 5'- GAGCTTGCTCCTTCCTTCTTAC -3' |
| *Cd31* | 5'- TGCTCACCATGCTGCTCTAT -3' | 5'- CAGTTTCTGCCCATTCGATAC -3' |
| *Enos* | 5'- TGACCCTCACCGATACAACA -3' | 5'- TCTGGCCTTCTGCTCATTTT -3' |
| *Epas1* | 5'- ATCCTTGTTCAAGCCACACC -3' | 5'- TTGCCATAGGTCGAGGATTC -3' |
| *Gata2* | 5'- ACATCCTACCTACGCCAACG -3' | 5'- GTGGCTTCAGCCAGACTAGG-3' |
| *Ang1* | 5'- TGGAGGAGGATGGACAGTAATA-3' | 5'- AGCTCGATCCTCAGCATGTACT-3' |
| *Tie2* | 5'- AGCGAGTAGACCATGCGAGT -3' | 5'- TTCGGCATCAGACACAAGAG -3' |

**Table 4. Fetal lung autopsy specimens from babies with CDH and human fetal lung specimens from elected terminations.**

| Case # | Age (Gestational weeks) | Description |
|---|---|---|
| 1 | 19 | Left-sided CDH, pulmonary hypoplasia |
| 2 | 20 | Left-sided CDH, pulmonary hypoplasia |
| 3 | 26 | Absence of left diaphragm, pulmonary hypoplasia |
| 4 | 27 | Left-sided CDH, pulmonary hypoplasia, intrauterine growth restriction, stillborn |
| 5 | 19 | Elected termination of pregnancy |
| 6 | 18 | Elected termination of pregnancy |
| 7 | 18.2 | Elected termination of pregnancy |
| 8 | 19 | Elected termination of pregnancy |
| 9 | 19.4 | Elected termination of pregnancy |
| 10 | 18 | Elected termination of pregnancy |
| 11 | 18 | Elected termination of pregnancy |
| 12 | 18.6 | Elected termination of pregnancy |
